# Molecular modeling of Proteinase-Activated Receptor 1 in complex with Thrombin Receptor Activator Peptide 6

**DOI:** 10.1101/2024.06.05.597686

**Authors:** E Reboul, D. El Hamaoui, S. Pasquali, P. Gaussem, E. Rossi, A. Taly

## Abstract

The protease-activated receptor 1 (PAR1) and its activator thrombin receptor activator peptide 6 (TRAP6) play crucial roles in various physiological and pathological processes, including hemostasis, thrombosis, and cancer progression. Although the interaction between PAR1 and TRAP6 has been heavily studied using experimental technique such as mutagenesis, structural data remains scarce due to the technical hardship of studying membrane proteins such as PAR1.

In this study, we employed an integrative modeling approach to elucidate the structure of the PAR1-TRAP6 complex. Leveraging state-of-the-art AI-based protein modeling tools, including AlphaFold2 and ESMFOLD, we integrated HADDOCK, a physics-based method to refine predictions. Overall, the predicted structures are in good agreement with the experimental data available in the literature. Our model unveiled a new T-shaped pi-stacking interaction between TRAP6’s F2 and PAR1’s Y360.

The integrative modeling approach combining the predictions of the deep learning model with a physics-based method proves to be an interesting strategy for solving challenging membrane protein structures with high confidence. Our model of the PAR1-TRAP6 complex will be an interesting starting point for further investigation of the activation of PAR1 by TRAP6.

## 1. Introduction

Protease-activated receptors (PARs) constitute a family of G protein-coupled receptors (GPCRs) present on various cell types, including those associated with the blood vessel wall (such as endothelial cells, fibroblasts, myocytes) and blood components (like platelets, neutrophils, macrophages) (1). Structurally, they feature an N-terminal exodomain joined to a seven-transmembrane helix (TM) bundle structure connected by intracellular and extracellular transmembrane loops (ICL, ECL), along with a C-terminal cytoplasmic tail (2). PARs become activated through N-terminus cleavage by plasma proteases at defined protease-specific sites, resulting in the generation of a tethered ligand. This latter binds intramolecularly to the ligand-binding site, leading to a conformational change that initiates a cascade of intracellular signaling events (3). Protease-activated receptor 1 (PAR1), was first identified as the major thrombin receptor on platelets (4). Thrombin is a crucial and multifaceted serine protease that promotes rapid and strong platelet aggregation, fibrin formation and amplification as well as negative regulation of the coagulation pathway. In platelets, PAR1 activation serves as a pivotal trigger for various responses. Thrombin-mediated PAR1 activation induces a change in platelet shape and promotes aggregation, a critical step in hemostasis. Moreover, PAR1 signaling contributes to the amplification of platelet activation, leading to the formation of stable blood clots. Understanding the intricate interplay between PAR1 and platelet function is essential for unraveling the complexities of thrombotic disorders (5; 6). Beyond its role in platelets, PAR1 plays a significant role in endothelial cells lining blood vessels. Activation of PAR1 in endothelial cells elicits diverse responses, including the release of vasoactive molecules and modulation of vascular tone. Additionally, PAR1 activation is involved in the regulation of endothelial permeability, a process crucial for maintaining vascular homeostasis. Positioned at the crossroads of platelet activation and endothelial cell function, PAR1 plays a crucial role in hemostasis and vascular biology. Imbalances in PAR1 activity are well-characterized contributors to disease pathophysiology and have been proposed to contribute in vascular disease, tumor metastasis, inflammatory disorders, and cardiovascular disease, highlighting its therapeutic potential (7; 8).

Unraveling a deeper understanding of PAR1 signaling in both humans and mice not only enhances our fundamental understanding of these processes but also opens avenues for developing targeted interventions for cardiovascular and thrombotic disorders. Investigation into PAR1 in both human and murine models has provided valuable insights into its physiological and pathological roles. The common features of PAR1 across these systems emphasize the significance of murine studies in understanding human biology. Moreover, the pharmacological targeting of PAR1 has emerged as a potential therapeutic strategy for conditions associated with aberrant coagulation and vascular dysfunction (9).

Importantly, the anti-PAR1 vorapaxar was approved in the USA in 2014 for use in addition to standard antiplatelet therapy for secondary prevention in patients with a history of myocardial infarction or symptomatic peripheral artery disease; however, its net clinical benefit remains uncertain (10; 11; 12). Given the bleeding risk, it has not gained EMA approval. Its benefit must be weighed against the increase in bleeding events and it is contraindicated in patients with a history of stroke, TIA or intracranial hemorrhage. Its efficacy as a sole antiplatelet therapy is unknown and its use in the context of ACS in addition to standard antiplatelet agents showed no significant benefit (11). New molecules directed against PAR-1 such as pepducine are under development. To limit the bleeding risk, anti PAR-4 are also currently in phase 1 and 2 of development.

The structure of PAR1 has been extensively studied, yielding a comprehensive understanding of its architecture. The distinctive feature of PAR1 is its activation mechanism through proteolysis of the N-terminal region by serine proteases between Arg-41 and Ser-42, to expose a new N-terminus that carries the SFLLRN sequence (Ser-42 to Asn-47). Through X-ray crystallography studies and electron microscopy, researchers have elucidated the three-dimensional structure of PAR1. These approaches have allowed researchers to visualize the specific arrangement of amino acids and domains in the receptor, providing a detailed understanding of its molecular architecture. It is important to note that the structure of PAR1 can exhibit conformational variations depending on its active or inactive state. This structural flexibility is crucial for understanding how the receptor responds to proteolysis and initiates a cascade of intracellular signals (13).

Thrombin Receptor Activator Peptide 6 (TRAP6), also called Selective Protease-Activated Receptor 1 (PAR-1) agonist, is a synthetic hexapeptide designed to mimic the N-terminal fragment of PAR-1 generated by thrombin cleavage (Ser-42 to Asn-47). Remarkably, it exhibits exclusive full agonist properties for PAR-1 activation. This synthetic peptide serves as a pharmacological tool to imitate PAR-1 activation without the necessity for proteolytic cleavage, thereby inducing platelet aggregation (14). Therefore, TRAP6 offers a cost-effective solution for conducting in-vitro studies on platelet activation in hematology.

Researchers use TRAP6 in experiments to specifically activate PAR1, allowing for the study of downstream signaling pathways and cellular responses associated with this activation. This synthetic peptide offers a controlled and reproducible way to explore the effects of PAR1 activation, circumventing the reliance on endogenous proteases such as thrombin. By doing so, it provides valuable insights into the role of PAR1 in various physiological and pathological processes. Thus, the binding of TRAP6 to PAR1 is a crucial molecular interaction, contributing significantly to our understanding of the intricate signaling cascades associated with PAR1 activation in platelets, endothelial cells, and other cell types (15).

## 2. Methods

### 2.1 sequence selection

The original sequence of 425 amino-acid was obtained from uniprot (16), the uniprot id is P25116. The modelling was limited to the mature receptor (matPAR1) which does not cary the peptide signal : 1MGPRRLLL-VAACFSLCGPLLS21 (17). The sequence of TRAP6 used is the following 1SFLLRN6.

### 2.2 ab initio prediction

All scripts for deep learning models are obtained via ColabFold v1.5.5 (18), which is a package that offers a user-friendly interface to run deep learning models for protein’s 3D predictions such as ESMFOLD (19) and AlphaFold 2 (AF2) multimer v3 (20) or AF2 ptm, both version using MMseqs2 (21) for Multi Sequence Alignement (MSA), and then executed on Google Colab(22).

For AlphaFold 2 multimer with MMseq2, the use of template was enabled for some predictions (see results). The MSA query was made with MMseq2 using unique reference environment. The alignment was made with paired sequence of the same species and additional unpaired sequence was used for each chain. We used the greedy strategy for pair alignment which pairs any taxonomically matching subsets instead of the complete strategy, where all sequences have to match in one line.

The number of recycles depends on the type of prediction. For multimer, i.e matPAR1/TRAP6 complex, the number of recycles was set to 3 with a tolerance threshold for early recycle stop set to 0. For monomer, i.e mat-PAR1, the number of recycles was set to 20 with early tolerance threshold set to 0.5.

For sampling, the maximum msa parameter was set to auto and the dropout option was not used.

All predictions were performed without the minimisation option, and a custom minimisation protocol was used instead (see below).

### 2.3. minimisation protocol

The ab initio prediction were minimize with a 1000 step steepest descent with a 0.02 Angstrom step size on Linux 64-bit UCSF chimera 1.17.3 (23). The hydrogen bonds were considered for minimisation and the protonation state was inferred based on the name of residues. The charge of standard residues was based on forcefield AMBER ffs14b (24).

### 2.4. Docking refinement with Haddock

To refine the poses of the matPAR1/TRAP6 complex, we used HAD-DOCK webserver (25; 26). TRAP6 was classified as a peptide and PAR1 as a protein receptor. Haddock uses Ambiguous Interaction Restraints (AIRs), which are user defined based usually on available experimental data. The AIR are defined using active residues and passives residue. Active residues are residues identified to be involved in the interaction between the receptor and ligand (peptide) and solvent accessible residues (residues with less 15% of surface accessible to solvent are automatically removed). Passive residues are are all solvent accessible surface neighbors of active residues.

In our case, all the residues of TRAP6 were defined as interactive. For matPAR1, the active residues were defined based upon a ligand-based pocket definition. Based on the minimized ab initio prediction 3D structure of the matPAR1/TRAP6, all PAR1 residues within a 5 angstrom radius of TRAP6 were defined as active residues. For the rest of the HADDOCK experience, the default parameters were used.

### 2.5. structure analysis

The structural exploration and visualisation of structure were performed using Linux 64-bit UCSF chimera 1.17.3 (23).

The interaction between PAR1 and TRAP6 was analysed using Protein Ligand Profiler (PliP) (27) websever and pre-built Singularity image for Linux version 2.3.0. Singularity version 3.8.6 (28) was used to run the PliP image. For the webserver and Singularity image, the inter/peptide mode was used to define TRAP6 as the ligand and PAR1 as the receptor.

### 2.6. modelling pipeline

We implemented a 4-step pipeline to model PAR1/TRAP6 complex. The first step is the creation of a template for AF2 using ESMFOLD, as illustrated in Figure 1 (a). The sequence of matPAR1 is given as an input to ESMFOLD. The 3D structure predicted by ESMFOLD is minimized using the protocol described above. The second step is the prediction of mat-PAR1’s 3D structure as shown in Figure 1 (b). The 3D structure prediction generated in the first step is used as a template for AF2 multimer with the same input sequence and also using the same minimisation protocol.

**Figure 1:**
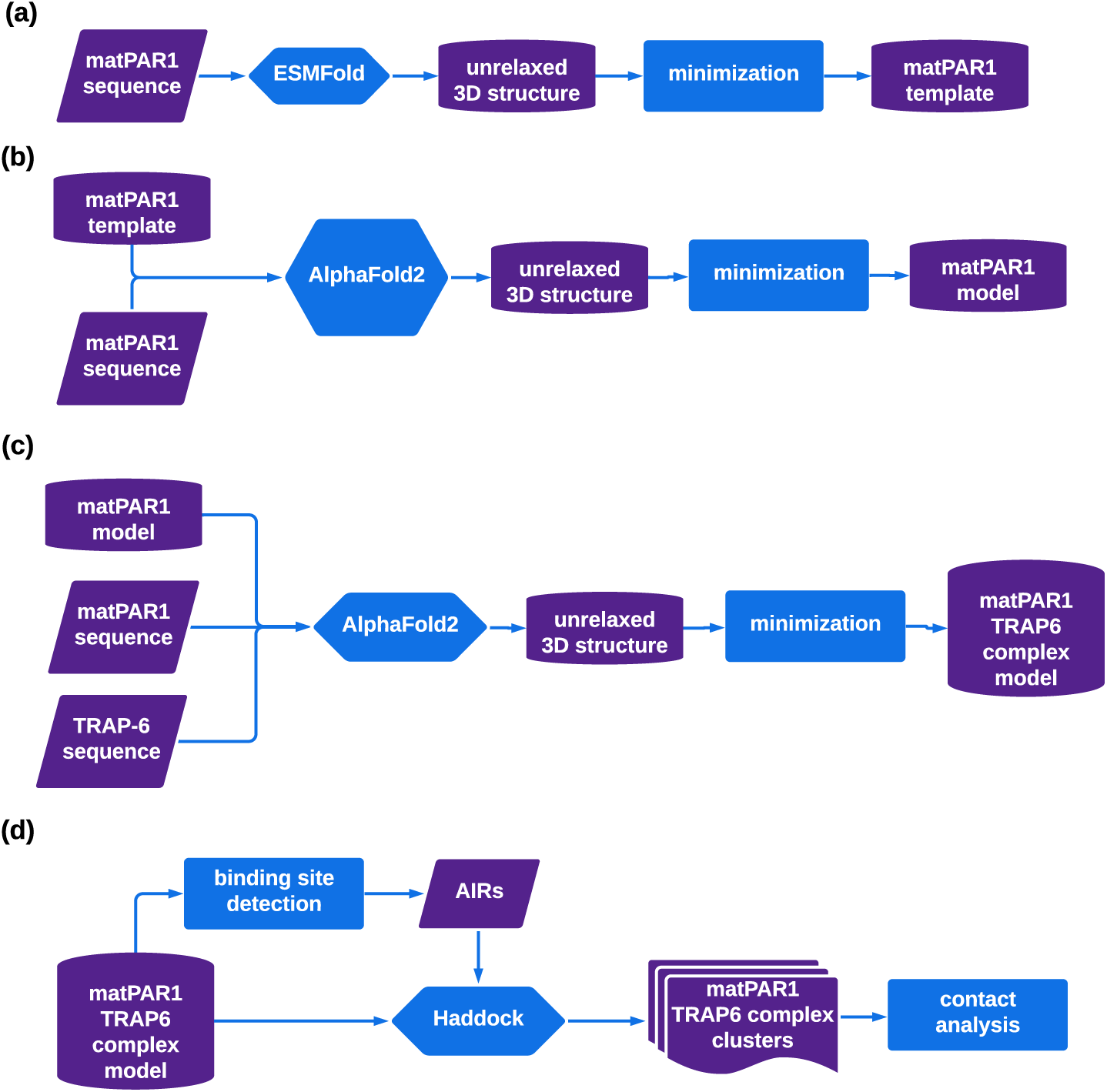
Modelling PAR1/TRAP6 complex : (a) the sequence of matPAR1 (22-425) is given as input to ESMFOLD, the unrelaxed output structure is minimized using our custom protocol. (b) the steps of the previous step are repeated with AF2 with MMseq2 for Multi sequence alignement. The output of the previous step is used as template structure for AF2. (c) AF2 mutlimer is used to generate PAR1/TRAP6 complex with output of the previous step as template for AF2 multimer (d) The prediction of PAR1/TRAP6 complex from the previous step is used to define AIRS and is used as the basis for Haddock docking refinement, all results are analysed using PLiP.

The third step is the prediction of the matPAR1 TRAP6 complex as shown in Figure 1 (c). The template structure is the final prediction of PAR1’s 3D structure and the sequence of TRAP6 (1SFRLLN6) is also given as an input. The sequence are differentiated using the symbol “:” as separator.

The last step is the TRAP6 pose refinement using Haddock as shown in Figure 1 (d). The active residues of AIR are defined using the AA in a 5 Angstrom radius of the TRAP6 ligand in the PAR1/TRAP6 complex in the previous step. All the clusters of structure generated by HADDOCK were analysed using PliP as detailled in subsection 2.5.

#### 3. Results

### 3.1. Modelling matPAR1

The only high resolution structure (2.2 angstrom) available for PAR1 in the PDB is the entry 3VW7 where PAR1 forms a complex with vorapaxar a specific inhibitor of PAR1 (29). The protein is actually a PAR1-T4L mutant with multiple modifications to make the crystallogenesis process easier. The putative N-ter disordered region was truncated by the insertion of a TEV protease after P85. The two N-linked glycosylation site in ExtraCellular Loop 2 (ECL2) have been removed by the following mutation : N250G and N259S. The C-ter domain was truncated after residue S395. A T4 lysozyme residues 2–161 were inserted in the third intracellular loop (ICL3) in place of the residue V302.

Therefore the first part of our experiment is dedicated to the modelling of the native PAR1 using Deep learning techniques. In our First attempt, we used AF2 with MM2seq and without template to predict matPAR1 alone.

As shown in Figure 2 (a) The TransMembrane helixes (TM) are well defined with predominately high to very high confidence in predicted structure, associated with pLDDT higher than 70. However the N-ter and C-ter have low pLDDT. For the N-ter, the first ninety AA are expected to be disordered with no particular secondary structure. Therefore, it is unlikely, that we can make a significant improvement on the prediction for the N-ter domain. However the C-ter of other GCPR of family A presents an alpha helix (30) which might indicate that the results can be improved for C-ter domain of matPAR1. The low quality of this portion of the model can probably be attributed to a low sequence coverage (Figure 3).

**Figure 2:**
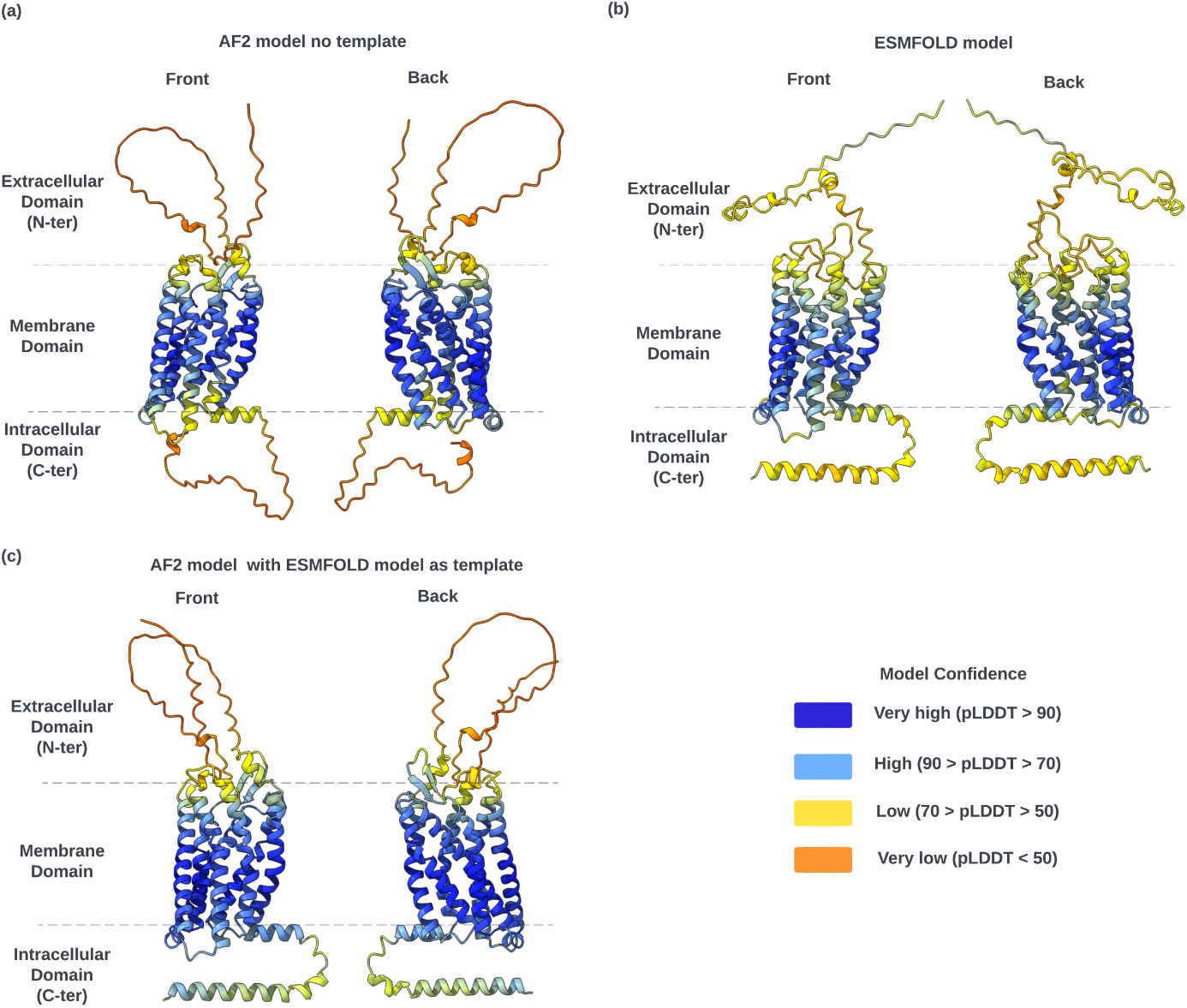
(a) front (left) and back (right) view of AF2 best ranked model prepared without a template. (b) front (left) and back (right) view of ESMFOLD model. (c) (a) front (left) and back (right) view of AF2 best ranked model prepared without the minimized structure of ESMFOLD model.

**Figure 3:**
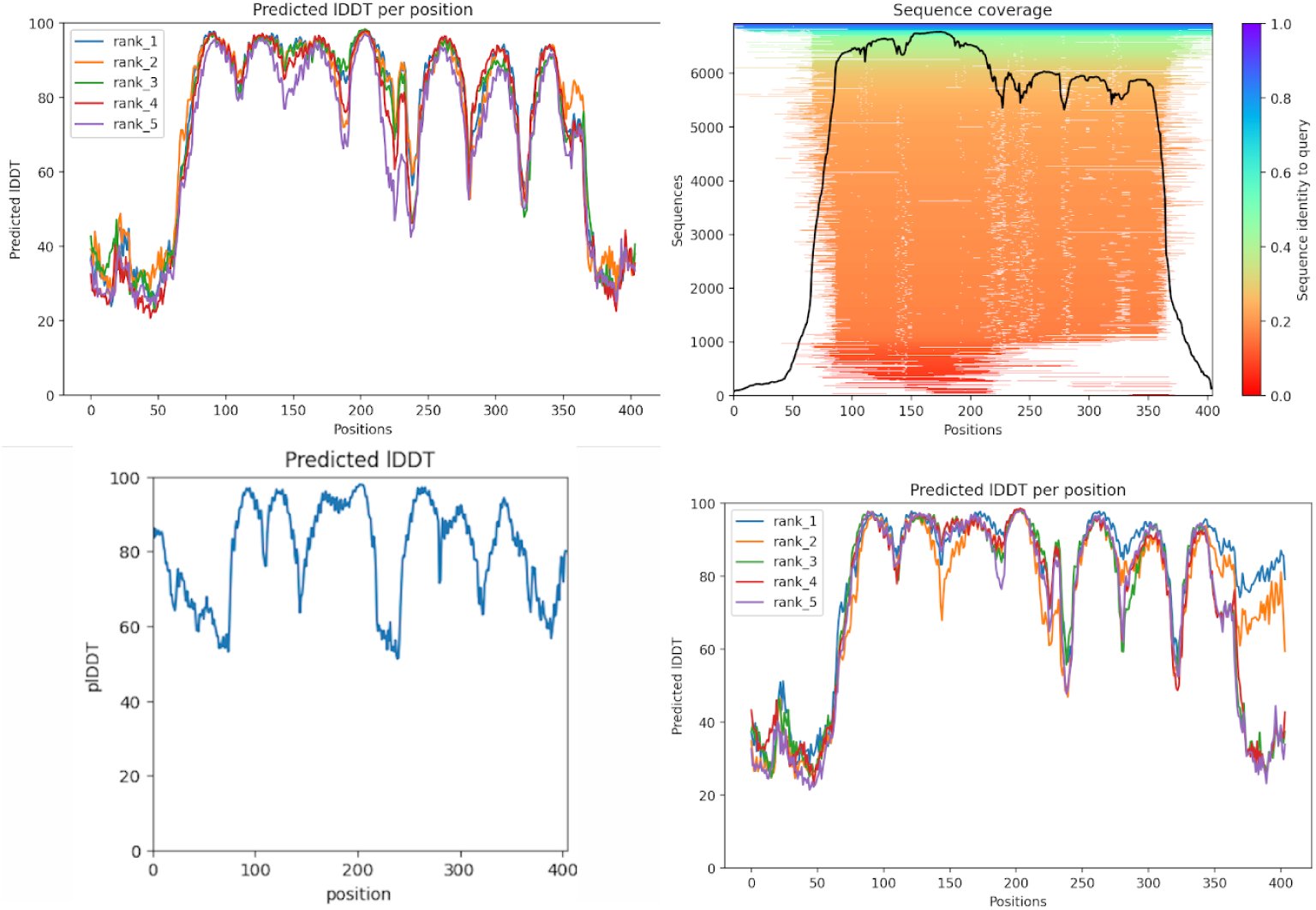
Improvement of AF2 models using a template generated with ESMFOLD. Top left: pLDDT score for five AF2 models generated without template. top right: Sequence coverage. Bottom left: pLDDT of ESMFold models. Bottom right: pLDDT scores of AF2 models using an ESMfold model as a template.

ESMFold doesn’t require a MSA to make a 3D structure prediction. We therefore reasoned that it could be able to propose a better model for the C-terminus of PAR1. Indeed, the models generated with ESMFold show better pLDDT score for the C-ter domain (Figure 3) and a very well defined helix as shown in Figure 2 (b).

This model was then used as a template for a new modelling with AF2. Two of the five models generated have a large pLDDT score for the C-ter domain, higher than those obtained previously by both ESMFolD and AF2 (Figure 3). The best ranked model of AF2 with template is shown in Figure 2 (c).

An interesting thing to notice is that AF2 seems to have taken the best of both ESMFOLD model and AF2 without template prediction in terms of secondary structure definition. This is shown with the reduced dip in pLDDT around the 150 and 225 residues shown in the bottom right corner of Figure 3.

### 3.2. Modelling matPAR1/TRAP6

Using the final PAR1 model described in subsection 3.1 as template, the 3D structure of matPAR1/TRAP6 complex was predicted using AF2 multimer as illustrated in the Figure 4 (a) and (b). The models generated by AF2 multimer put TRAP6 inside the binding pocket identified by the PAR1-vorapaxar complex (29) (not shown).

**Figure 4:**
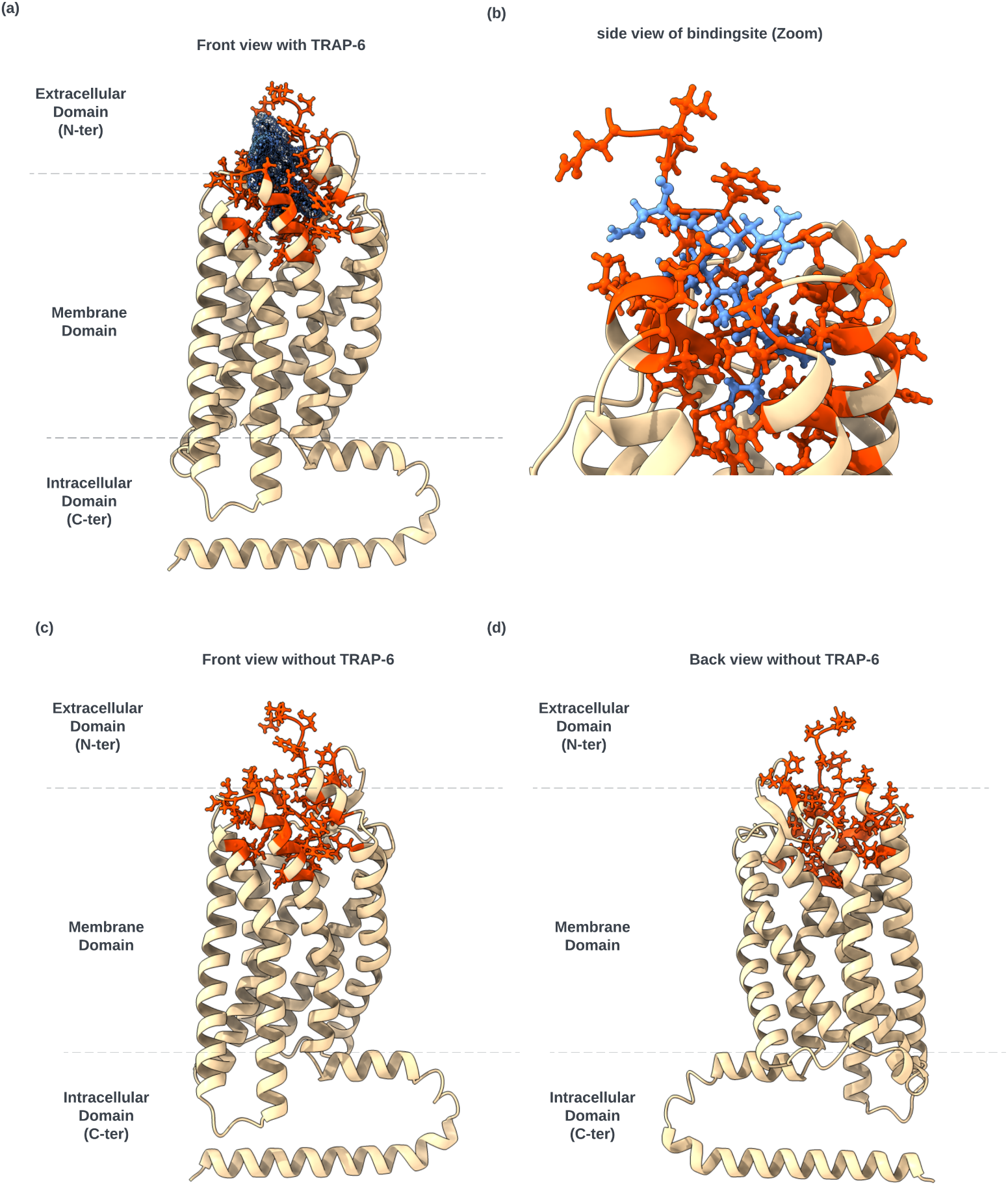
Structure of the first ranked PAR1-TRAP6 complex obtained with AF2 multimer using MM2seq. TRAP6(cornflower blue) and side chain of residues (orange red) witin a 5 Angstrom radius are represented as balls and sticks. The rest of the matPAR1 structure is represented as cartoon. For clarity sake, the disordered region of the N-ter domain (1-82) is not shown. (a) front view of the PAR1-TRAP6 complex, The surface drawn as a cloud of dot is superimposed on the structure of TRAP6 (b) Zoomed in side view of the binding pocket with TRAP6 in it (c)/(d) front and back view of PAR1 without TRAP6

However, a Force Distance based Atomic Force Microscopy (FD-based AFM) study (31) shows that the binding of the tethered ligand obtained by the cleavage by thrombin has two meta stable states : a shallow binding and a deep binding. The current finding seems to indicate that AF2 multimer with default sampling option generates only the shallow binding.

Therefore we decided to refine the docking using HADDOCK, a physics-based methods of docking. We defined of the Ambiguous Interaction Restriction (AIR) to consider the residues of PAR1 within a 5 Angstrom radius of TRAP6 in the previous step to be active residues as shown in Figure 4 (c) and (d). All residues of TRAP6 are defined as active residues.

To select the deep and shallow binding poses in Haddock we implemented different criteria :

- a well define secondary structure
- contact between residues identified by mutagenesis or other experimental methods
- position In/On the binding site for Deep/shallow binding respectively
- a difference in predicted affinity between shallow and deep binding

First, mutagenesis studies confirmed that there is a specific interaction between PAR1’s E264 et E260 and the TRAP6’s R5 where a switch between this two amino-acid produce viable binding of the tethered ligand to PAR1. We also looked for residues whose mutations are associated with a significant decrease in PAR1 activity, namely: L96, D256 and E347. Another critical information known from the literature is the mutated TRAP6 peptide S**A**LLRN T(F2A) has a singular drop in affinity for PAR1. This entail that the key interaction between TRAP6 phenylalanine is not a Van Der Waals interaction but rather a Pi-stacking interaction or Pi-cation interaction.

The main criteria to discriminate between shallow binding and deep binding is the position of the backbone of TRAP6. For the deep binding the backbone should be inside the binding side of PAR1 and for the shallow binding the backbone should be at the opening of the PAR1 binding pocket. Another consideration should be that HADDOCK’s scoring function should reflect the relative difference of binding affinity of both poses, i.e the predicted affinity by haddock for the deep binding pose should be higher than for the shallow one.

We found poses of TRAP6 consistent with a shallow binding pose and a deep binding pose, as depicted in Figure 5 (a) and (b) respectively. The deep binding pose show that the head of TRAP6, in particular its unique phenylalanine, is inserted deep into PAR1 binding site as shown in Figure 5 (a) and (c). The Tail of the TRAP6 is at the opening of the binding site. Interestingly a parallel beta-sheet is formed between the TRAP6 ligand and the Extra Cellular Loop 2 (ECL2).

**Figure 5:**
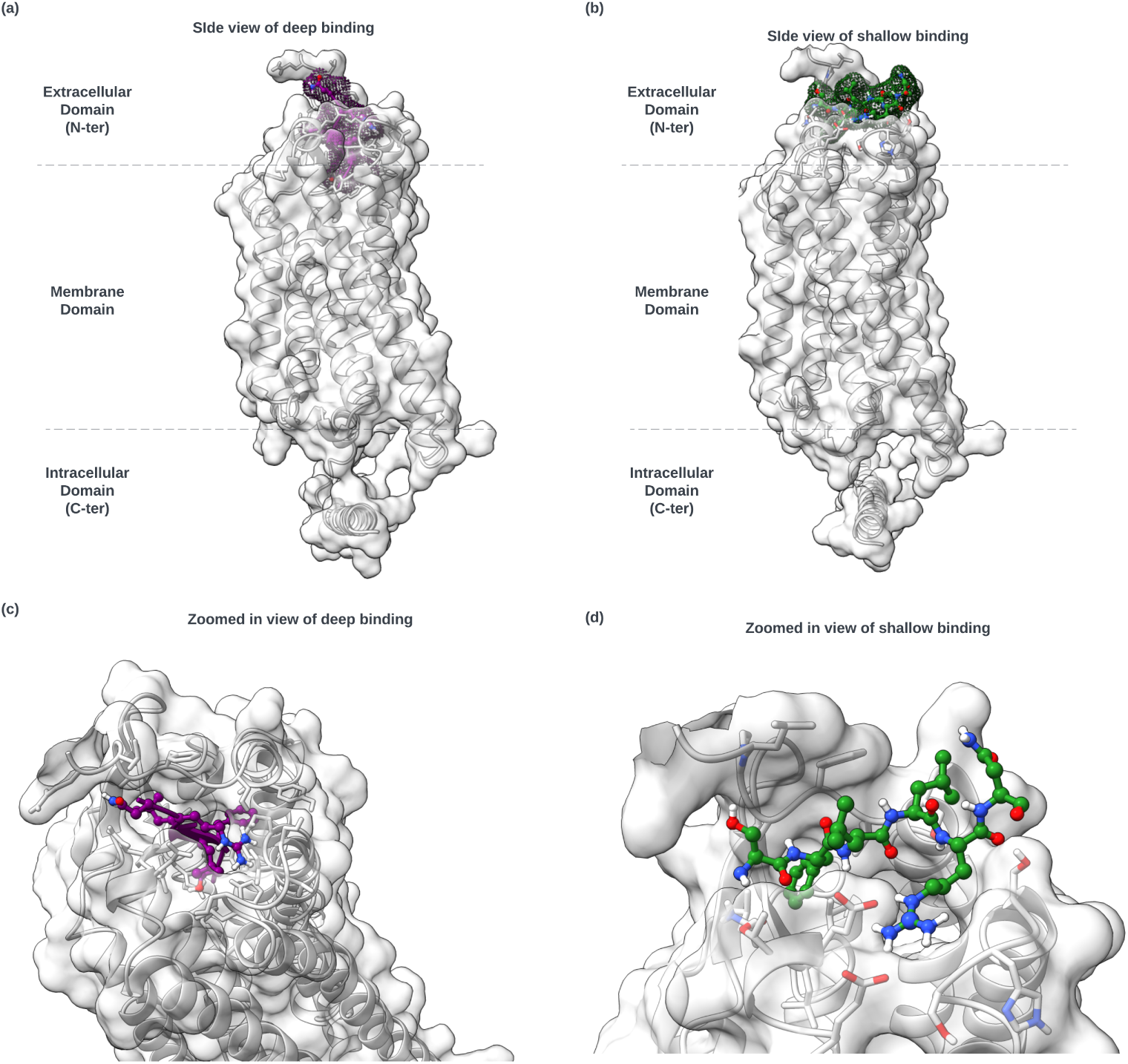
Structures of the PAR1-TRAP6 complex obtained after docking refinement using Haddock. (a) and (c) represent the deep binding, with the side view for (a) and a zoom in view on the binding pocket for (c). (b) and (d) represent the shallow binding.

In contrast, for the shallow binding TRAP6 is placed at the opening of PAR1 binding pocket. TRAP6’s F2 is placed in a small hydrophobic pocket formed by the outer layer of ECL2 and the end of the N-termini disorder region, as illustrated in Figure 5 (b) and (d).

We took a deeper dive into the interaction between PAR1 and TRAP6 for both the shallow and deep binding pose with PLiP (27). The detail for every binding pose is available in the Supplementary information. The deep binding pose is the second pose of the third cluster and the shallow binding pose is the the third pose of the sixth cluster. We can note that HADDOCK classify the cluster by energy, i.e cluster 3 has a lower predicted energy which is consistent with the discrepancy in term of affinity between the shallow and the deep binding pose.

For the deep binding pose we found two salt bridges between TRAP6 Arginine (R5) and PAR1’s E260 and E347. TRAP6 L4 is placed in a small hydrophobic pocket formed by the end of the N-ter dirsordered domain and ECL2 by the following residues sidechains : A86, I87 and V257. TRAP6’s L3 is placed in a small hydrophobic pocket made by chain of Q260’s *γ* carbon, L263 side-chain, L340 side chain and T346’s gamma carbon. There is a perpendicular (T-shape) pi stacking between TRAP6 F2 and Y350, coupled with binding to otherwise hydrophobic pocket. This hydrophobic pocket is composed of the following residues : Y95, A92, E347 *γ* carbon and V257. TRAP6’s S1 share hydrogen bond with H340 via the oxygen (acceptor) of the amid group, and with Y337 via the oxygen of its side-chain (donnor).

There is an extensive hydrogen bond network, that goes beyond the interaction with TRAP6’s S1, with a total of 15 between PAR1 and TRAP6 deep binding pose. Part of the hydrogen bond network allows for the formation of 2 small parrallel beta sheet between TRAP6’s F2, L3, L4 and PAR1’s ECL2 residues : V257, L258, N259. The parallel beta sheets are maintained by 4 hydrogen bonds between the backbones of TRAP6 and PAR1 : F2-D256, F2-V257, L4-L258 and L4-Q260.

For the shallow binding pose, we found overall less interaction between TRAP6 and PAR1. The most interacting TRAP6’s residues are F2 and R5, which is coherent with the mutagenesis studies in the literature. TRAP6’s F2 is inserted into a pocket formed by the en of the N-ter disordered domain and a part of ECL2. This pocket is formed by the following PAR1’s sidechain of residues : A86, I88, I244, V257. TRAP6’s R5 take part in two salt bridges with E260 and E264.

Regarding the rest of the interactions, they all are Hydrogen bonds, 7 in total. TRAP6’s residues with the most hydrogen bond interaction is S1 with 3 hydrogen bonds. TRAP6’s S1 share a hydrogen bond from it side chain hydroxyl group (donor) with the oxygen of PAR1’s P85 (acceptor). The other two originate from the unbounded nitrogen of S1’s backbone. The two acceptors of those hydrogen bonds are T261 and N259. The rest of the hydrogen bond are between:

- TRAP6’s F2 backbone oxygen (acceptor) and PAR1’s F87 backbone nitrogen (donor).
- TRAP6’s L3 backbone nitrogen (donor) and PAR1’s E260 charged sidechain carboyxlic acid (acceptor).
- TRAP6’s R5 backbone oxygen (acceptor) and PAR1’s S344 sidechain hydroxyl group (donor).
- TRAP6’s R5 sidechain guanidinine group (donor) and PAR1’s S341 backbone oxygen (acceptor).

**Figure 6:**
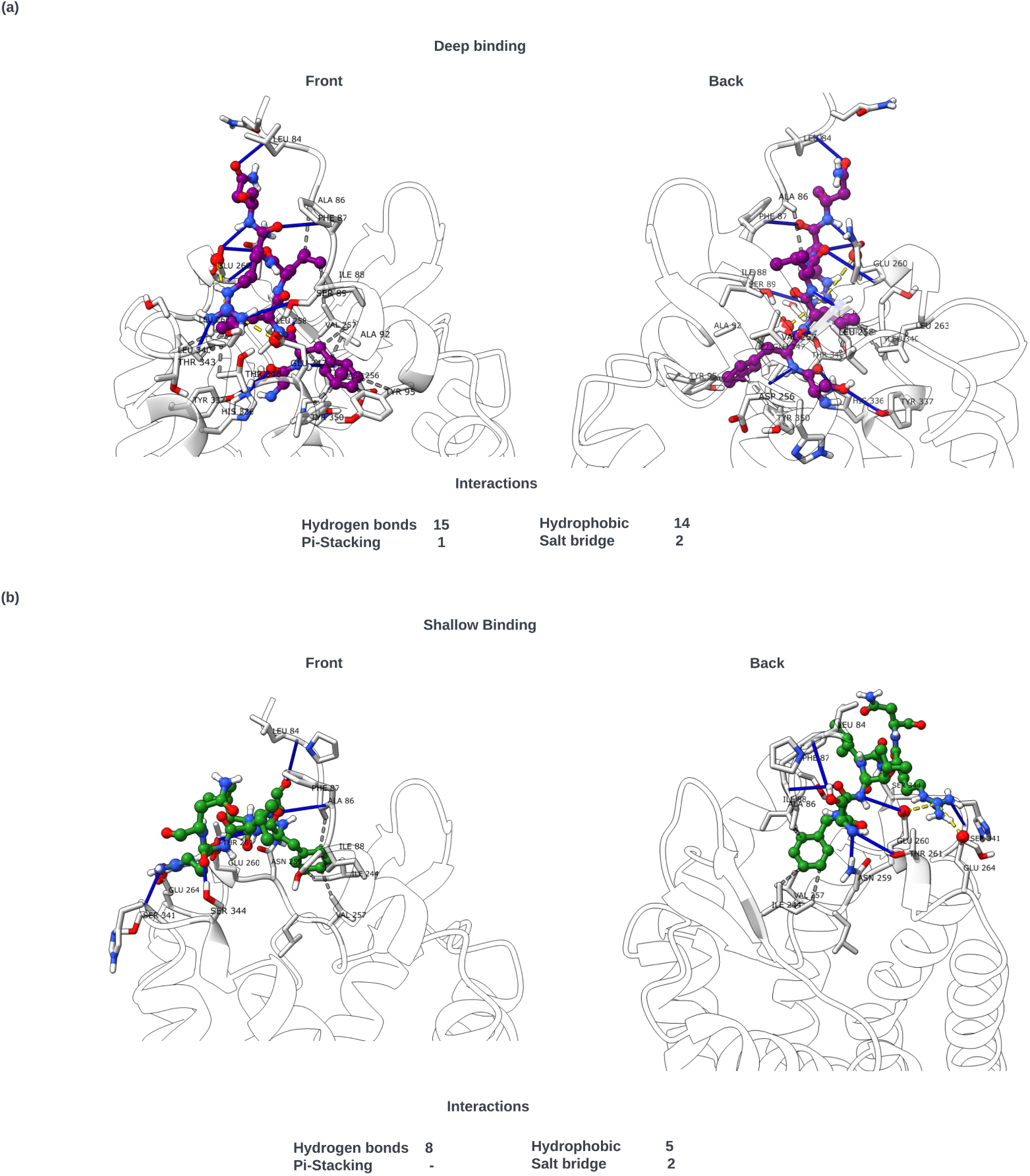
Interactions between PAR1 (light grey) and TRAP6 (Deep binding pose in Purple and green in cluster B) detected by the interaction analysis tool PLiP (27) for the two cluster. Interacting PAR1’s AA were label using the 3 letter code and their residue number (in black). For each complex hydrogen bonds (solid blue lines), *π*-stacking (dashed green line), hydrophobic contacts (dark gray dashed lines) and salt bridges (yellow dashed lines) are highlighted. Below each structure we report the total number of each kind of interaction detected.

### 3.3. Membrane integration and post-translational modification modelling

At last we used Chimera to edit the orientation of the sidechain of PAR1’s C387 and C388 using the rotamer editing tool based *Shapalov et al* rotamer library (32). We selected the rotamer which best pointed toward the membrane. We then used CHARMM-GUI (33; 34; 35) membrane builder (36; 37; 38; 39) to add the Palmitoylation to PAR1’s C387 and 388. Note that the N-glycolisation of N250 and N259 and other PAR1’s residues is possible, however the lack of clear experimental data on the N-glycolisation profile of those position doesn’t allow their confident modelling.

PAR1 was oriented in the membrane using the PPM 2.0 webserver (40) and inserted in a symmetrical bilayer membrane composed of 230 POPC per leaflet. A water padding of 22.5 angström was added to both the extracel-lullar and intracellular part of the membrane. The system was neutralised using potassium and chlore ions. As depicted in Figure 7, we can see that PAR1 fits in the membrane with its two palmitoyl anchors inserted in the lower leaflet of the bilayer membrane.

**Figure 7:**
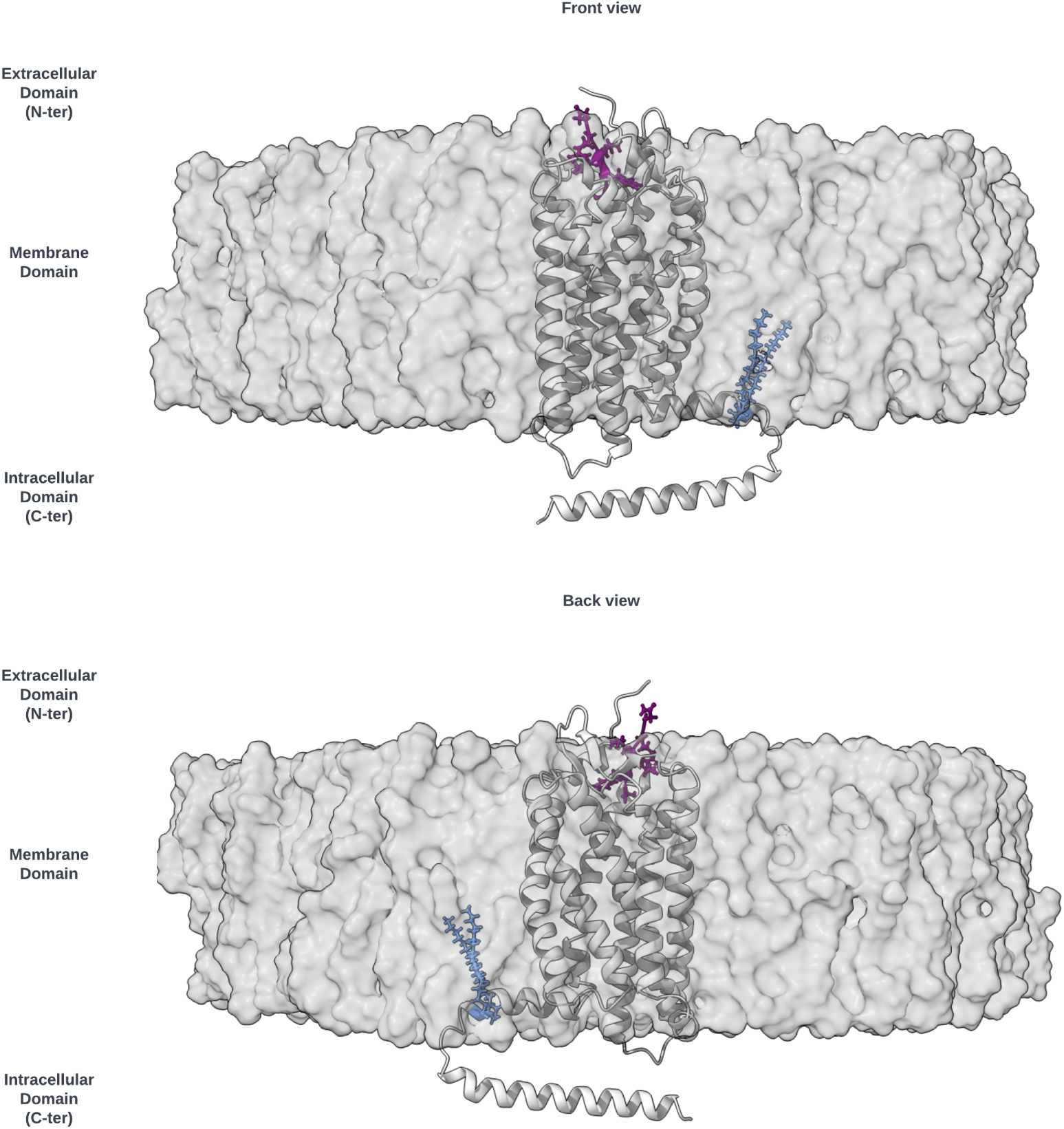
PAR1 TRAP6 complex in a 100 % POPC membrane, depicted as the grey semi-transparent surface. PAR1 is depicted with light grey cartoon structure. TRAP6 in its deep binding pose is represented as a purple ball stick structure. At last PAR1’s C387 and C388 side chain are represented also as ball stick colored in cornflower blue. The solvent molecule, i.e water and ions are not represented

## 4. Discussion

The binding of TRAP6 to PAR1 is a crucial molecular interaction, contributing significantly to our understanding of the intricate signaling cascades associated with PAR1 activation in platelets, endothelial cells, and other cell types (15). Thus, the aims of our work is to allow for a better understanding of the mechanism in which PAR1 is activated by TRAP6.

### 4.1. PAR1-TRAP6 complex modelling

To that end, we first tried to model PAR1 alone using AF2. Unfortunately, C-terminus domain was poorly resolved probably due to poor sequence coverage in the MSA. To overcome this challenge we used a pure NLP model for 3D prediction, ESMFOLD. The prediction made by ESMFOLD proposed a well folded C-termini region. It was used as a template for AF2 to make a prediction that combine the strength of both AF2 and ESMFOLD. The TRAP6 PAR1 complex was initially model using the prediction obtained in the previous step as a template for AF2 multimer.

In the literature 2 binding modes are described for the endogenous tethered peptide to PAR1: a shallow binding and a deep binding (31). AF2 was able to find poses of TRAP6 that are consistent to a deep binding. However, with the given input parameter, AF2 multimer was unable to find the shallow binding mode. Using HADDOCK allowed to overcome this limitation.

### 4.2. Interaction of TRAP6 with PAR1

Our analysis of the interaction of TRAP6 with PAR1 has confirmed that both a shallow and deep binding are observed with our pipeline.

The well established theory of salt bridges between PAR1 E260, E264 residues and TRAP6 R5 has been verified for the shallow binding pose, and only E260 for the deep binding pose. PAR1’s E347 has been identified in the literature as critical for PAR1 activation, and is confirmed by it’s salt bridge with TRAP6’s R5 when TRAP6 is in the deep binding pose.

The importance of the extra cellular loop 2 (ECL2) has also been confirmed with the formation of small parallel beta sheets between PAR1 and TRAP6 via hydrogen bonds formed between respective backbones when TRAP6 is in the deep binding pose. It also participate in the formation of a small hydrophobic pocket in which shallow binding TRAP6-s F2 and deep binding TRAP6’s L4 interact.

Alanine scan performed in the literature showed a steep decrease in affinity for TRAP6 mutation F2A which pointed to an non-hydrophobic inter-action to be the key interaction between TRAP6’s F2 and PAR1. Our modelling results suggest that there is a strong perpendicular (T-shaped) pi-stacking interaction between TRAP6’s F2 and PAR1’s Y350. Moreover, although the direct interaction between PAR1’s D256 and TRAP6 are limited, an analysis of the hydrogen bond network shows D256 to be an anchor point connected to Y350, Y161, Y162 and K158. This result suggests that D256 is a key residue into maintaining the proper orientation of the Y350 and in the transduction signal.

Hopefully, our results will allow for a better understanding of the mechanism in which PAR1 is activated by TRAP6. The specific targeting of PAR1 Y350 via pi-stacking might reveal a great way to emulate a TRAP6 agonist effect.

### 4.3. Modelling with a combination of modelling tools

The combination of ESMFOLD and AF2 multimer with MM2seq for MSA shows promising results. The model of an orphan sequence (with low coverage in MSA), which shows limited results with AF2, was improved using a template generated by ESMFold. Indeed, the resulting model seems to benefit from the combination of the respective strengths of both methods, at least as can be assessed from the pLDDT overall score, which is better for the fusion model than for the separated models. Furthermore, the improvement of the model with Haddock then opens up the possibility to take into account additional aspects like Post-translational modifications and ligand binding. This observation is coherent with recent observation that the combination of deep-learning based methods and molecular dynamics can be fruitful (41; 42; 43).

### 4.4. Therapeutic implications of discoveries

The knowledge of the TRAP6 interaction with PAR-1 paves the way for developing new therapeutic molecules using TRAP6 mechanism of action as a model/template. If PAR-1 inhibitors have been marketed for a long time, peptides that activate PAR-1 have not been developed so far, while fibrinogen-associated thrombin has been used as a hemostatic patch a long time ago. This probably comes from the numerous preclinical data published at the end of the last century showing a disparate effect of TRAP6 systemic administration, varying according the species and the dose administered. For instance, mouse platelet main thrombin receptor is PAR4. Experiments on rats showed that their platelets are resistant to TRAP, whereas rat vasculature is highly sensitive to TRAP. These data suggest that the thrombin receptors on rat vasculature may be similar to those on human platelets, whereas the receptors and/or the coupling mechanisms in rat platelets are different from human platelets (44). As a result, when infused to rats, TRAP induced an endothelium-dependent vasodilatation that leads to a decrease in arterial blood pressure, this effect being at least in part mediated by the endothelial release of NO (45).

In guinea-pig and in primates, platelets are responsive to TRAP, contrary to other animal species. As an example, when administered to guinea-pig, contrary to rats, TRAP9 induced a bronchoconstriction through a mechanism involving platelets since depleting animals of circulating platelets prevented this bronchoconstriction (46). Because of the pleiotropic effect of thrombin and of TRAP administered intravenously, their administration in humans is challenging. To date, only thrombin has an indication in human therapy restricted to local administration as hemostatic patch.

Despite the inability to use TRAP6 as a therapeutic agent due to its multiorgan effect particularly on vessel wall and platelets, understanding its interaction with the PAR1 receptor is useful for applications in fields such as:

i. Studying Platelet Activation PAR-1 is the major thrombin receptor on platelets, and TRAP6 mimics the action of thrombin without its enzymatic activity, making TRAP6 a useful tool to study PAR-1 platelet responses
ii. Research on Coagulation Pathways By activating PAR-1, we can explore the signaling mechanisms and downstream effects involved in coagulation and thrombosis
iii. Drug Development and Testing TRAP6 is used ex vivo in preclinical and clinical studies to evaluate the efficacy of antiplatelet drugs, with no necessity to obtain washed platelets. Indeed, by inducing platelet activation without clotting in PRP samples, we can simply assess how well new therapeutic agents inhibit this process, which is crucial for developing treatments for cardiovascular diseases. TRAP6 is also recommended in clinical practice as one of the platelet agonists to be used for exploration of platelet functions in bleeding syndromes.
iv. Studying Inflammatory Responses Beyond its role in hemostasis, PAR-1 is involved in inflammatory responses. TRAP6 can be used to investigate the role of PAR-1 in various inflammatory diseases, helping to elucidate potential therapeutic targets for conditions like atherosclerosis.

